# *Colletotrichum* spp. from soybean cause disease on lupin but can also induce plant growth-promoting effects

**DOI:** 10.1101/2020.06.19.160853

**Authors:** Louisa Wirtz, Nelson Sidnei Massola Júnior, Renata Rebellato Linhares de Castro, Brigitte Ruge-Wehling, Ulrich Schaffrath, Marco Loehrer

## Abstract

Protein crop plants such as soybean and lupin attract increasing attention because of their potential use as forage, green manure or for the production of oil and protein for human consumption. While soybean production only recently gained more importance in Germany and within the whole EU in frame of protein strategies, lupin production already is well established in Germany. The cultivation of lupins is impeded by the hemibiotrophic ascomycete *Colletotrichum lupini*, the causing agent of anthracnose disease. Worldwide, soybean is also a host for a variety of *Colletotrichum* species, but so far this seems not to be the case in Germany. Cross-virulence between lupin and soybean infecting isolates is a potential threat, especially taking into consideration the overlap of possible soybean and lupine growing areas in Germany. To address this question, we systematically investigated the interaction of different *Colletotrichum* species isolated from soybean in Brazil on actual German soybean and lupin plant cultivars. Conversely, we tested the interaction of a German field isolate of *C. lupini* with soybean. Under controlled conditions, *Colletotrichum* species from soybean and lupin were able to cross-infect the other host plant with varying degrees of virulence, thus underpinning the potential risk of increased anthracnose diseases in the future. Interestingly, we observed a pronounced plant growth-promoting effect for some host-pathogen combinations which might open the route to the use of beneficial biological agents in lupine and soybean production.

## Introduction

In view of the steadily growing world population and the increasing demand for sustainable food production, it becomes clear that dietary protein cannot be provided through animal products alone. Soybean is the primary source for non-animal protein and the most cultivated protein-crop plant on a global scale. In Europe soybean and soybean-derived products are imported for the most part (Lucas et al. 2015). Because European consumers shift their purchase behavior more and more to non-GMO products from sustainable production soybean imports from abroad are under debate. Hence, the independent production of protein-crop plants in Germany becomes increasingly attractive, and lupin represents an appealing alternative to soybean (Lucas et al. 2015; Talhinhas et al. 2016).

In the past decades, lupin production was negatively affected by the hemibiotrophic ascomycete *Colletotrichum lupini*, causing anthracnose disease (Nirenberg et al. 2002; Talhinhas et al. 2016; Pecchia et al. 2019). The influence of anthracnose epidemics was severe, partially due to focusing on breeding of low-alkaloid containing white lupin in the past, which is especially susceptible to *C. lupini* (Berger et al. 2012). Breeding for resistance of lupin against *C. lupini* is slowly progressing, but still the impact of the disease on production success is severe and alternative plant protection methods are needed (Fischer et al. 2015; Jacob et al. 2017).

Currently, soybean cultivation in Germany is mostly affected by the fungal diseases *Sclerotinia sclerotiorum* causing stem rot (“white mold”) and the disease complex *Diaporthe spp./Phomopsis* spp, causing seed and stem blight (Ploper and Backman 1992; Murphy-Bokern et al. 2017; Tian et al. 2017). As *Colletotrichum* is a significant problem in worldwide soybean production and because of the increasing production, it is only a matter of time until anthracnose will become a severe problem on soybean in Germany.

Recently the causal agents of anthracnose on soybean were systematically investigated in Brazil, describing *C. truncatum* as the major anthracnose-causing species but also identifying *C. plurivorum* as a novel species capable of causing anthracnose (Rogério et al. 2017, 2019; Barbieri et al. 2017; Dias et al. 2018; Damm et al. 2019).

In this study, we performed inoculation experiments with *Colletotrichum* species isolated from soybean and lupin in Brazil and Germany, respectively, on plant cultivars used in Germany to evaluate the risk of cross-infection in a scenario of increased legume production. Results from this study shall contribute to planning of current and future plant breeding goals and open the road to novel plant protection strategies.

## Material and Methods

### Plant material and growth conditions

Seeds of *Glycine max* cv. Abelina (not inoculated with rhizobia) and *Lupinus angustifolius* cv. Lila Baer were kindly provided by I.G. Pflanzenzucht GmbH, Munich, Germany. Plants were grown in a climate chamber (day/night cycle: 16 h light, 350 μmol m^-2^ s^-1^ PAR, 24 °C, 65 % RH; 8 h dark, 20 °C, 80 % RH) in ED73 substrate (Balster Einheitserdewerk GmbH, Fröndenberg, Germany).

### Fungal isolates and culture conditions

The German field isolate of *Colletotrichum lupini* BBA70358 (=CBS 109222) is deposited at the CBS collection of the Westerdijk Fungal Biodiversity Institute (Utrecht, The Netherlands) and was donated to B. Ruge-Wehling by H. I. Nirenberg. *C. truncatum* isolates (see table 1) were provided by N.S. Massola Jr. and (University of São Paulo, ESALQ, Department of Plant Pathology and Nematology, Piracicaba/SP, Brazil) stored at the culture collection of Plant Pathology Department of ‘Luiz de Queiroz’ College of Agriculture, University of São Paulo, Brazil (Rogério et al. 2017). Isolates of *C. plurivorum* (table 1) were also provided by N.S. Massola Jr. and stored at the culture collection of Plant Pathology Department of ‘Luiz de Queiroz’ College of Agriculture, University of São Paulo, Brazil (Barbieri et al. 2017). Richard O’Connell (BIOGER, INRA, Paris, France) kindly provided *C. higginsianum* IMI 349063. All fungi were propagated on PDA medium (potato extract glucose agar, Carl Roth GmbH + Co. KG, Karlsruhe, Germany). The fungal cultures were incubated at 25°C with a 12 h day/night cycle under cool white fluorescent lamps. Every 7-10 days fungal isolates were transferred to fresh PDA plates. To prevent degeneration of isolates (i. e. reduced sporulation), at regular intervals (every three months) material from stocks was used to start a new culture.

**Table 1.** fungal isolates used in this study.

| species | strain | isolated from |
| --- | --- | --- |
| *Colletotrichum lupini* | BBA70358 (=CBS109222) | lupin |
| *Colletotrichum truncatum* | LFN150 | soybean |
| *Colletotrichum truncatum* | LFN151 | soybean |
| *Colletotrichum truncatum* | LFN152 | soybean |
| *Colletotrichum truncatum* | LFN153 | soybean |
| *Colletotrichum truncatum* | LFN154 | soybean |
| *Colletotrichum plurivorum* | LFN0007 | soybean |
| *Colletotrichum plurivorum* | LFN0008 | soybean |
| *Colletotrichum plurivorum* | LFN0010 (=URM7540) | soybean |
| *Colletotrichum plurivorum* | LFN0018 (=URM7541) | soybean |
| *Colletotrichum plurivorum* | LFN0019 (=URM7542) | soybean |
| *Colletotrichum plurivorum* | LFN0023 | soybean |
| *Colletotrichum higginsianum* | IMI349063 („063A“) | Arabidopsis |

### Hypocotyl assay

The protocol for the toothpick inoculation assay was adapted from Scandiani et al. (2011). To prepare hypocotyls of soybean and lupin, seeds were placed on wet filter paper and incubated in the dark. Seeds that germinated after 2-3 days were planted in ED73 soil and placed in a climate chamber as described above. The hypocotyls were ready for the assay at the plant growth stage of formation of the first leaf. Sterile toothpick tips (1.0—1.5 cm) were placed around a growing fungal colony on PDA plates and used for hypocotyl inoculation when fully overgrown by fungal mycelium. The inoculated toothpick tips were pushed into hypocotyls using sterile forceps. After seven days lesion length was measured using a Leica MZ125 stereo-microscope equipped with a digital JVC KYF 750 camera using Diskus software (Technisches Büro Hilgers, Köngiswinter, Germany).

### Seed inoculation assay

Soybean and lupin seeds were surface sterilized by washing in 70 % EtOH for 1’, followed by 3x rinsing in sterile distilled water. Afterwards, the seeds were washed in 1 % sodium hypochlorite solution again followed by 3x rinsing in sterile distilled water. Seeds were dried under sterile conditions for 24 h. For this assay, PDA-mannitol agar plates, adjusted to an osmotic potential of 1 MPa by adding 74,69 g*l^-1^ D-mannitol, were prepared, following the method described in Scandiani et al. (2011). Fungal cultures were grown on PDA-mannitol plates until the colonies spread all over the plates. Surface sterilized seeds were placed on those plates and incubated for 48 h under growth conditions described for fungal cultures above. The inoculated seeds were planted in ED73 substrate, and seedling growth was monitored. The mock control consisted of surface-sterilized seeds, that were incubated on PDA-mannitol plates without fungus.

## Results

### Cross-infection assays on soybean and lupin hypocotyls

Some plant pathogenic fungi from the genus *Colletotrichum* are known to have a broad host range and, in addition, their lifestyles can range from necrotrophy to endophytic behavior (Talhinhas et al. 2016; De Silva et al. 2017). In this study, we firstly investigated *Colletotrichum* species originating from soybean or lupin for their ability to cross-infect both hosts. The infection process of *Colletotrichum* species on host plants is mediated by a specific infection structure called appressorium. This specialized cell is melanized and facilitate a direct penetration of the host cuticle and cell wall by weakening the plant tissue with the help of secreted lytic enzymes in combination with physical pressure driving the entry of the infection hypha (Bechinger 1999; Küster et al. 2008; Ludwig et al. 2014; Loehrer et al. 2014). To assess the principal ability of the *Colletotrichum* isolates under investigation to colonize a host plant irrespectively of appressorium formation, we firstly employed an infection assay based on fungus-inoculated toothpicks that are directly placed into the seedling’s hypocotyls. In case of successful colonization, lesions are formed at the site wounding caused by the toothpicks, and lesion length in comparison to the mock-inoculated control served as a measure for virulence (Fig. 1a). Different field isolates of *C. truncatum* caused the largest lesions on hypocotyls of the German soybean cultivar Abelina (maturity group 000). Mean lesion lengths of the *C. truncatum* isolates LFN150, LFN152 and LFN154 were 5.1, 5.4 and 4.8 mm, respectively, and significantly different from the mean lesion length of 1,8 mm caused by the sterile, non-inoculated toothpick tip alone (Fig. 1a). Infection with isolates of *C. plurivorum* and *C. lupini*, as well as *C. higginsianum*, also resulted in lesions larger than the control but the test for statistical significance failed. For the experiments with lupin, two different plant species were chosen: *L. albus* (broad-leafed, white-flowering sweet lupin) and *L. angustifolius* (narrow-leafed, blue-flowering sweet lupin). *L. albus* cv. Amiga was most susceptible to *C. lupini* isolate BBA70358 (mean lesion length: 8.2 mm) (Fig.1b). Most interestingly, also the soybean-derived *C. truncatum* isolates LFN153, and LFN150 were able to cause lesion sizes (mean lesion length of 4.3 mm and 2.7 mm, respectively) which are significantly larger than the mock control but still not as large as those caused by *C. lupini*. Furthermore, the *C. plurivorum* isolate LFN0010, also derived from soybean, was able to colonize *L. albus* hypocotyls to a similar extent as the *C. truncatum* isolates (Fig.1b). A similar picture was obtained in the toothpick-inoculation experiments with *L. angustifolius* cv. Lila Baer. Again, the adapted *C. lupini* isolate BBA70385 caused the most prominent lesions (mean value of 11.46 mm lesion length – Fig. 1c). With mean lesion length ranging from 2.0 to 3.3 mm, the *C. truncatum* isolates LFN153, LFN152 and LFN154 caused smaller, but still significant, lesion lengths.

**Fig. 1.**
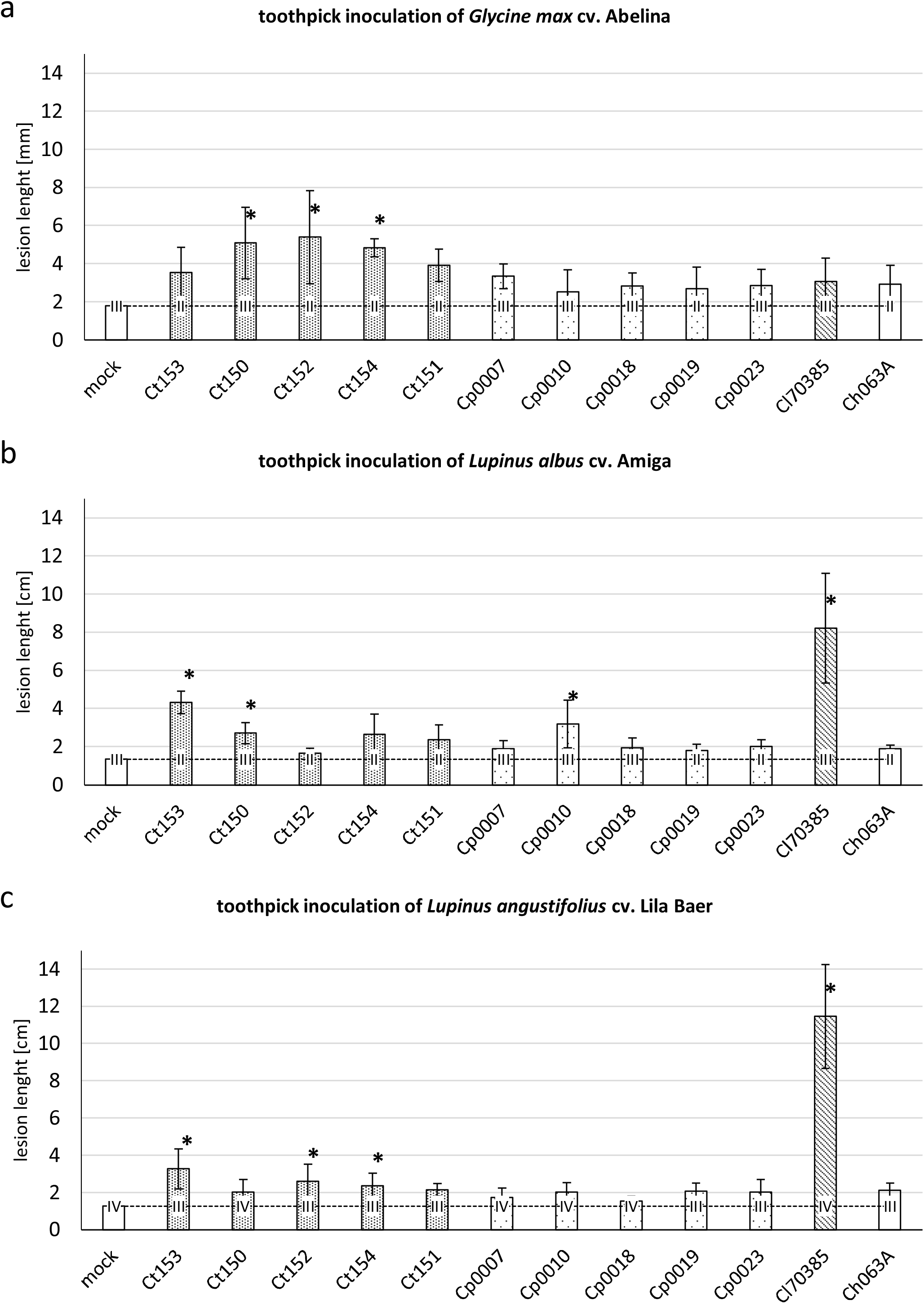
Results of lesion length measurements after inoculation of soybean and lupin hypocotyls. Hypocotyls of *G. max* cv. Abelina (a), *L. albus* cv. Amiga (b) and *L. angustifolius* cv. Lila Baer (c) were infected with *Colletotrichum*-inoculated toothpick tips and lesion lengths were measured after seven days. The abbreviations “*Ct*”, “*Cp*”, “*Cl*”, and “*Ch*” stand for the species *C. truncatum, C. plurivorum, C. lupini*, and *C. higginsianum*, respectively. Bars represent the mean value of two to four independent biological experiments with two plants each. Error bars show standard deviation. Asterisks indicate a statistically significant difference in comparison to mock control, determined by a one-way ANOVA analysis on ranks, followed by a Dunn’s posthoc test (P<0.05). In the graphs, the dashed line represents the mean value of the mock control.

### Cross-infection assays using inoculated seeds of soybean and lupin

To assess the cross-infectivity of different *Colletotrichum* isolates on soybean and lupin in an independent assay, we performed seed inoculation experiments. This assay is assumed to be closer to the natural way of plant infection because the fungus must infect the seedling without previous wounding. The same set of fungal isolates as in the toothpick-inoculation assay was tested on soybean cv. Abelina and narrow-leafed lupin cv. Lila Baer (Fig. 2). Because not all seeds made it to normal-looking seedlings, we chose a classification of five categories encompassing a range of non-emerged seedlings (0), seedlings which emerged but did not develop further (1), and seedlings which are smaller (2), equally (3) or better (4) developed in comparison to the mock control. On soybean, all *C. truncatum* isolates had severe effects on seedling emergence, in most cases inhibiting emergence. *C. plurivorum* isolates LFN0007, LFN0010, LFN0018, LFN0023, the *C. lupini* isolate, and the *C. higginsianum* isolate caused retardation in seedling development in comparison to mock-treated seeds one week after sowing.

**Fig. 2.**
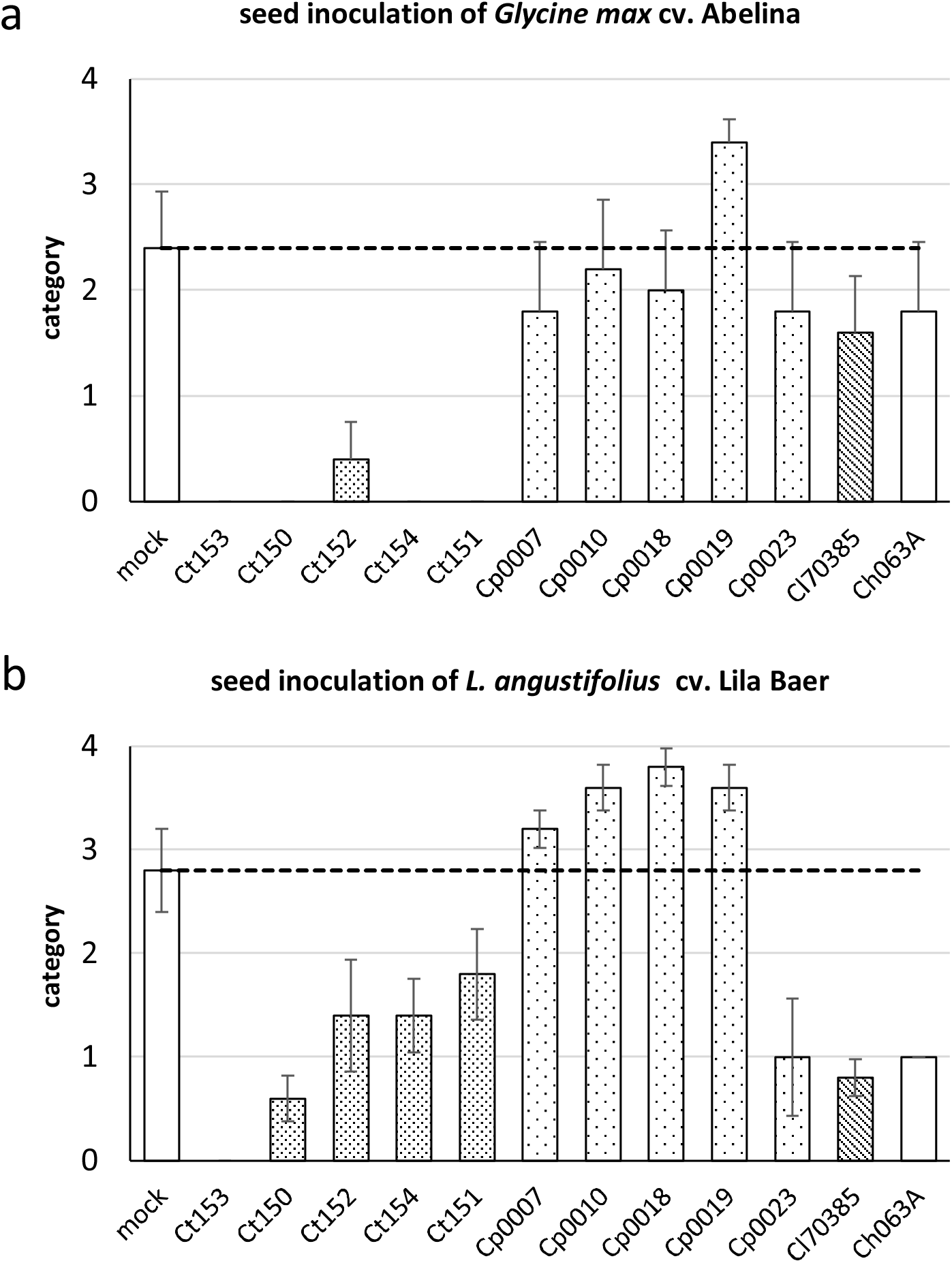
Rating of seedling emergence and development after inoculation of soybean and lupin seeds with *Colletotrichum* isolates. Seeds of *G. max* cv. Abelina (a) and *L. angustifolius* cv. Lila Baer plants (b) were inoculated with different *Colletotrichum* species and rated one week after sowing. The abbreviations “*Ct*”, “*Cp*”, “*Cl*”, and “*Ch*” stand for the species *C. truncatum, C. plurivorum, C. lupini*, and *C. higginsianum*, respectively. The following categories for rating were applied: 0 – seedling not emerged; 1 – seedling development arrested after emergence; 2 – slower or impaired seedling development in comparison to seedlings emerging from mock-inoculated seeds; 3 seedling development like mock; 4 – faster/better development in comparison to mock. Bars represent the mean value of five plants and error bars show standard deviation. In both graphs, the dashed line represents the mean value of the mock control.

Interestingly, seeds inoculated with *C. plurivorum* isolate LFN0019 developed better in comparison to non-inoculated seeds (Fig. 2a). This plant growth-promoting effect of *C. plurivorum* was even more prominent in the seed-inoculation experiment with *L. angustifolius* cv. Lila Baer (Fig. 2b). Here all but one isolate of *C. plurivorum* led to seedlings, that developed better than seeds from the non-inoculated control. In this experiment, the *C. plurivorum* isolate LFN0023 led to a similar reduction in seedling development as the *C. lupini* and the *C. higginsianum* isolate. All *C. truncatum* isolates clearly impaired lupin seedling development, and seed-inoculation with the *C. truncatum* isolate LFN153 completely prevented seedling emergence (Fig. 2b).

### Plant-growth promoting effect on lupin

The purpose of the seed inoculation experiments was to get an overview of the interactions of the whole range of fungal species and isolates in this study; therefore, they were based on single experiments with a lower number of plants. To support the observation regarding the plant growth-promoting effects, an independent biological experiment with *L. angustifolius* cv. Lila Baer and the *C. plurivorum* isolates was performed (Fig. 3). Instead of using our custom rating system, this time, the height of plants was measured, and the developmental stage was assessed according to a lupin-adapted BBCH scale. The plant developed from *C. plurivorum* inoculated seeds all were significantly taller than the plants originating from mock-treated seeds after seven weeks (Fig. 3a). The rating, according to a lupin-adapted BBCH scale showed that not only plant height was positively affected, but also the number of leaves (Fig. 3b). Whereas the tendency of an increased leaf number could be observed for all interactions with the fungal isolates, only interactions with *C. plurivorum* isolates LFN0010 and LFN0023 proved to be significantly different from mock-control. Figure 4 shows representative plants from this experiment seven weeks after sowing. The differences between plants of the non-inoculated control and those emerged from seeds treated with the fungus are easily visible (Fig. 4a). It seems that seedlings are not only taller after inoculation with *C. plurivorum*, rather the development as a whole is affected as some of the plants already progressed to flowering stage (Fig. 4b).

**Fig. 3.**
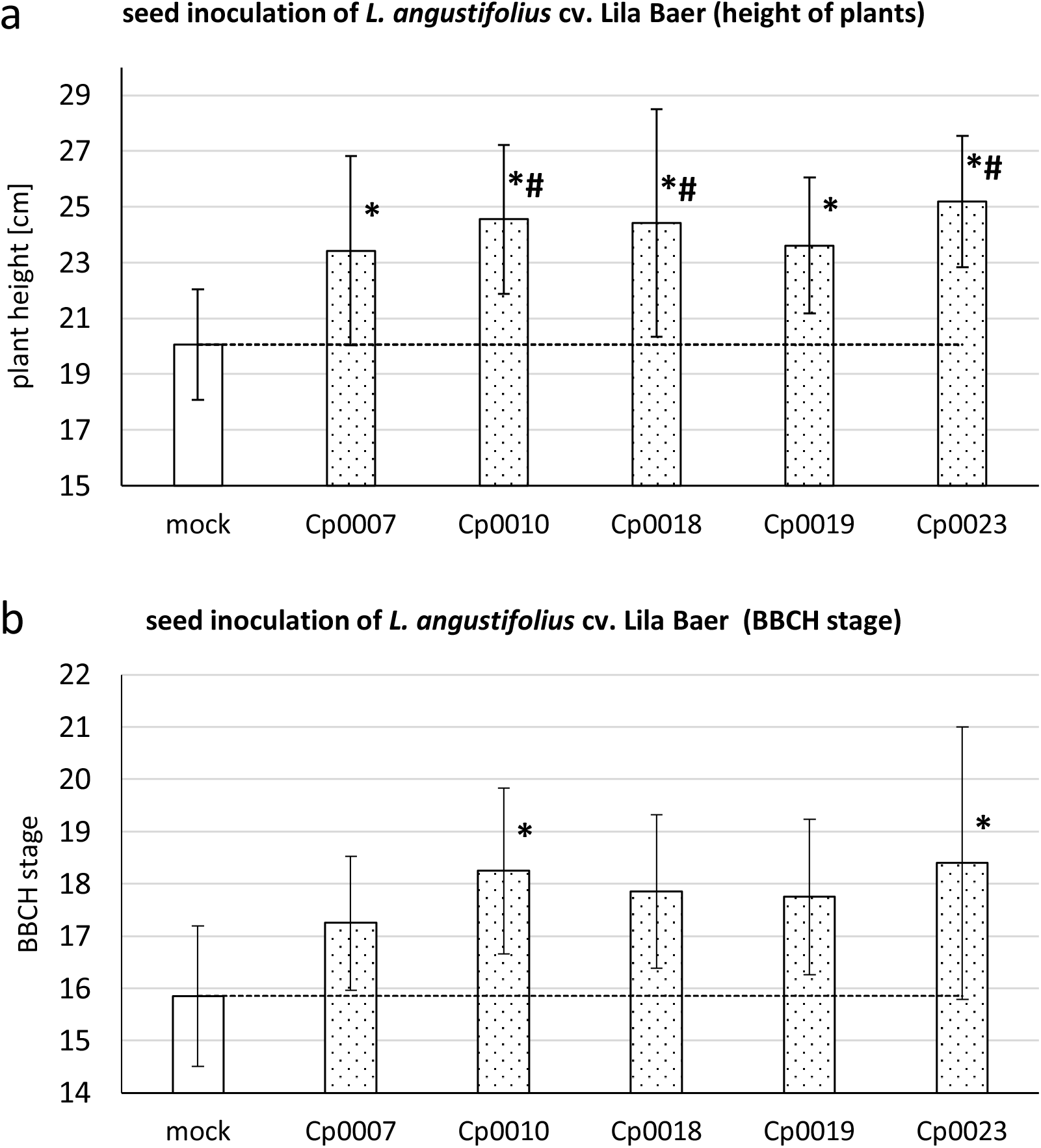
Plant-growth promoting effect of *C. plurivorum* inoculation on lupin development. In an independent experiment, heights of *L. angustifolius* cv. Lila Baer plants after inoculation of seeds with different *C. plurivorum* isolates were measured five weeks after sowing (a). The plants were also scored according to a lupin-adapted BBCH development scale (b). Bars represent the mean value of at least five plants and error bars show standard deviation. Statistical significance in relation to mock was tested with a one-way ANOVA analysis followed by Dunnett’s (P<0.05, number sign) or Holm-Sidak (P<0.05, asterisk) posthoc test (a). In b) test for statistical significance was performed by a one-way ANOVA analysis on ranks, followed by a Dunn’s posthoc test (P<0.05, asterisk). In both graphs, the dashed line represents the mean value of the mock control.

**Fig. 4.**
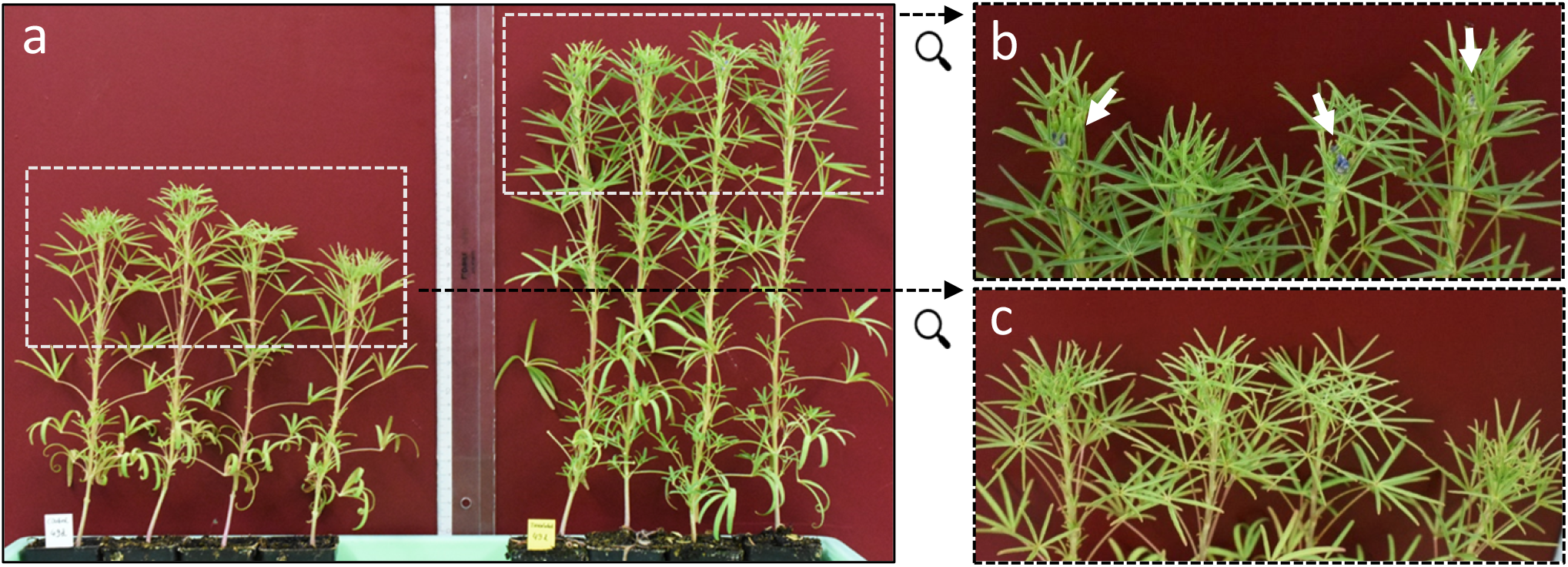
Plant-growth promoting effect of Colletotrichum *plurivorum* in seed inoculation assay. Comparison of plants from *L. angustifolius* cv. Lila Baer after treatment of seed without (a, left side) or with *C. plurivorum* isolate LFN0010 (a, right side) seven weeks after sowing. Part b) and c) of the figure are enlargements of the respective white dashed boxes in a), indicated by black arrows. White arrows in b) point to developing flowers. Pictures correspond to data shown in Fig. 3.

## Discussion

In this study, the interaction of *Colletotrichum spec*. isolates originating from soybean or lupin were tested for the first time on soybean and lupin cultivars actually used in Germany Since, to our knowledge, so far soybean anthracnose has not yet been systematically investigated on soybean in Germany, we made use of isolates of *C. truncatum* and *C. plurivorum* coming from Brazil (Rogério et al. 2017; Barbieri et al. 2017).

To get an overview of the virulence spectrum of the different fungal isolates on both host plants, at first, an assay with a simple read-out, namely measurement of lesion lengths, was performed. This method using toothpicks colonized by the fungi was already established and successfully applied for studying *Fusarium virguliforme* and *C. truncatum* and *C. plurivorum* on soybean and applied here for the first time to lupin hypocotyls (Scandiani et al. 2011; Rogério et al. 2017; Barbieri et al. 2017). This assay is very robust and the read-out in form of lesion length measurement is easily quantifiable. However, the inoculation method including wounding and is thus not closely related to the natural infection process. By wounding, the pre-formed resistance barriers of the cuticle and an intact epidermis are circumvented.

Differences in colonization, therefore, must be attributed to defense mechanisms active during the colonization of the plant in response to the invading fungus (Hutcheson 1998; Scandiani et al. 2011). On soybean cv. Abelina all *C. truncatum* colonized the hypocotyl beyond the wound-induced lesion, but only the isolates LFN150, LFN152, and LFN154 lead to significant large lesions (Fig 1a). The soybean-derived isolates of *C. plurivorum*, as well as *C. lupini* caused smaller lesions than the *C. truncatum* isolates. Remarkably, also *C. higginsianum*, known to naturally infect plants from the *Brassicaceae* family was able to invade the tissue to a certain extent albeit also not statistically significant (Fig. 1a). This rather unexpected result might be related to the artificial inoculation assay or could also be explained by the exceptional large host range of many species of the genus *Colletotrichum*, which possess the general ability to grow on a variety of plants as well as dying and dead tissue (Damm et al. 2012; De Silva et al. 2017).

In the toothpick assay, *C. lupini* caused the largest lesions on both lupin cultivars Amiga and Lila Baer (Fig. 1, b and c). The interactions of lupin with the other Colletotrichum isolates revealed a more differentiated picture between *L. albus* (Fig. 1, b) and *L. angustifolius* (Fig. 1, c), which might be attributed to differences in basal resistance of both lupin cultivars.

Since anthracnose of soybean and lupin is reported to be seed-born, we decided to also employ a seed infection assay to get closer to the natural infection process (Talhinhas et al. 2016; Pecchia et al. 2019). For this assay, seeds were co-incubated with the fungus on agar plates. The swelling of the seeds followed by germination, which quickly happens after placing seeds on agar-medium, would make this method inadequate We prevented swelling and germination by increasing the osmotic potential of the agar-medium, enabling a longer co-incubation phase for seed and fungus, resulting in very evenly inoculated seeds (Scandiani et al. 2011). Soybean seeds pre-treated this way with *C. truncatum* isolates only rarely developed seedlings, and again, also *C. lupini* and *C. higginsianum* showed a tendency to cause retarded seedling development (Fig 2, a). In the case of *L. angustifolius* cv. Lila Baer, the *C. truncatum* isolate LFN153 exhibited a stronger negative effect on seedling emergence, than *C. lupini* (Fig 2, b). Most interestingly, besides causing retardation of seedling development, some *C. plurivorum* interactions led to promotion of plant growth of both soybean and lupin seedlings (Fig. 2). These findings were confirmed in an independent experiment using *L. angustifolius* and the *C. plurivorum* isolates (Fig. 3). In addition to evaluation in the first experiment, the development of plants was monitored over a longer time span. After five weeks the plant growth-promoting effect could be observed for all interactions tested. This was not only manifested by increased plant height (Fig 3, a), but also by the advancement to more mature developmental stages as observed in comparison to plants growing from mock-treated seeds (Fig. 3, b). Thus, after seven weeks, some of the inoculated plants already progressed to flowering stage (Fig. 4.).

In contrast to the first experiment, the isolate LFN0023 also caused a plant growth-promoting effect. Our results underline the plasticity of the interaction of *Colletotrichum spec*. with different hosts which can range from pathogenicity to commensalism and mutualism. Maybe also external factors influence this balance, as a substrate poor in nutrients was used in the second experiment and plants were grown longer than in the first experiment. It is known, that *Colletotrichum* species exhibit a broad spectrum of different lifestyles ranging from endophytic growth to necrotrophic behavior and that dynamic transitions in lifestyle can happen depending on host plants and environmental factors (Hacquard et al. 2016; Hiruma et al. 2016; De Silva et al. 2017).

Our study revealed, that soybean-derived isolates of *Colletotrichum* spec. are capable of causing disease on lupin and that *C. lupini*, at least under laboratory conditions, can colonize soybean. Whether or not this may cause problems in legume production in Germany cannot be foreseen, but considering the overlap of areas where soybean and lupin production is possible, this might only be a matter of time (Fig. 5). In this study, we reported on a plant growth-promoting effect of fungi from the genus *Colletotrichum* for legumes; however, the underlying mechanism remains enigmatic. Future studies have to explore the mechanism and should address the question and whether the fungus can grow endophytically like, e.g., *Piriformospora indica* (syn. *Serendipita indica)* (Deshmukh et al. 2006; Zuccaro et al. 2011). It will be interesting to see whether this beneficial interaction might render plants more resistant to abiotic and biotic stresses. Ultimately these results could contribute to novel approaches in legume production.

**Fig. 5.**
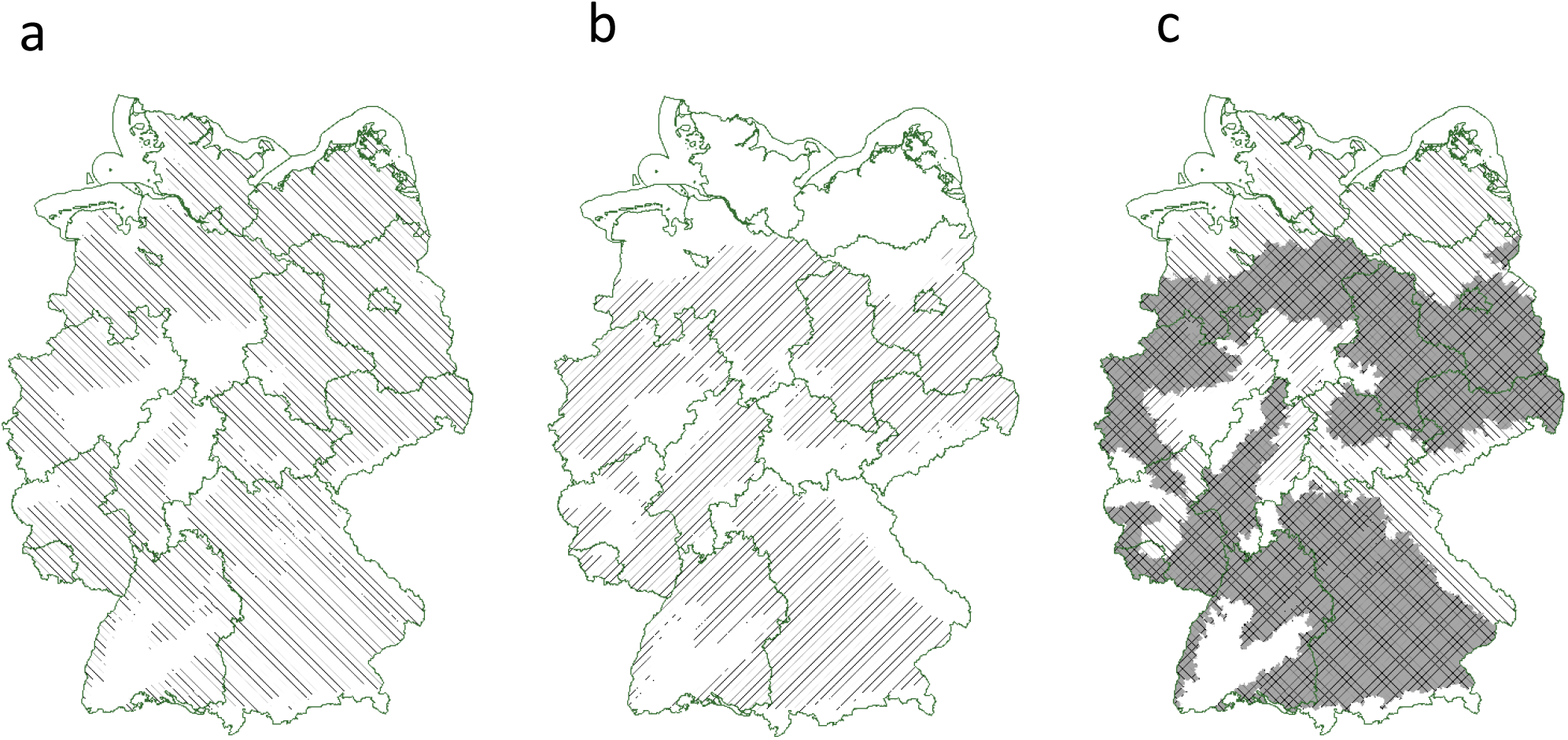
Comparison of potential cultivated area for narrow-leafed sweet lupin and soybean in Germany. Possible areas for cultivation of narrow-leafed sweet lupin (a, vertical hatching) and soybean (b, vertical hatching) based on soil-climate-areas are illustrated. An overlay of a) and b) shows the overlap of areas (c, gray-underlaid cross-hatching). Data for maps were obtained and modified from Julius Kühn-Institut (geoportal.julius-kuehn.de/index, accessed 08/2019) under license according to German GeoNutzV and from German Federal Agency for Cartography and Geodesy (http://www.bkg.bund.de, accessed 08/2019) under Data license Germany – attribution – Version 2.0 (dl-de/by-2-0).

## Author contributions

ML and US conceived and planned the experiments. Louisa Wirtz carried out the experiments, processed experimental data and drafted figures. ML analyzed the data, finalized the figures and drafted the manuscript. RRL and NSMJ conceived and adapted the inoculation assays on soybean, provided material and contributed to interpretation of results. BRW encouraged to study anthracnose on lupin, provided material and gave critical feedback on the results. All authors read and approved the final manuscript.

## Compliance with Ethical Standards

The authors declare that they have no conflict of interest. No experiments involving humans or animals have been conducted.

## Acknowledgments

The authors are grateful to Monika Hermanns for technical assistance. Many thanks to Richard O’Connell for kindly providing the *C. higginsianum* isolate.

## Notes

### Competing Interest Statement

The authors have declared no competing interest.

